# Crystal Structures of Ferritin Grown by the Microbatch Method in the Presence of Agarose and Electric Field Shows Enhanced Metal Binding

**DOI:** 10.1101/2021.09.30.462655

**Authors:** Aditya Seetharaman, Priyadharshine Ramesh Babu, Maham Ismail, Darcie J Miller, Vivian Stojanoff

## Abstract

Ferritin is an ubiquitous iron storage protein found in all kingdoms of life. Ferritin is essential for iron homeostasis and is involved in a wide range of physiological and pathological processes. Several structures of ferritin in complex with small molecules and metal ions have been reported. Here we report the crystal structures of Horse Spleen Ferritin, in which the crystals were grown by employing a novel approach adopting the microbatch experiments performed in the presence and absence of electric field using a 2% agarose pellet of CdSO_4_. We observed that 1) these structures contain increased number of Cd ions as compared to the crystallization of same protein by others using different methods. 2) The externally applied electric field reduced the number of nucleation and with fewer nucleation the size of the crystals increased. 3) There is no significant conformational change observed among these structures. 4) Irrespective of the externally applied electric field, this agarose microbatch crystallization method facilitates the retaining of increased number of bound metal ions with ferritin to mimic the possible *in vivo* environment.

## 1. Introduction

The intracellular protein ferritin is widely distributed and highly conserved throughout the animal and plant kingdoms[1]. Ferritins are among the most studied group of proteins involved in iron metabolism. There are two types of ferritins: L and H subunits. The major physiological functions of ferritins are to store and release iron in a controlled manner to protect against iron deficiency and iron overload. Ferritin also plays a key role in safeguarding cells from potentially toxic elements[2]. Iron is housed within ferritin nanocages, formed by the self-assembly of 24 ferritin subunits[3]. This unique assembly property of ferritin has been well explored to exploit the method of encapsulation and transport for other molecules such as anticancer drugs, bioactive molecules, and mineral metal ions [4]. Indeed, previous studies have reported ferritin in complex with several small molecules, including anesthetics and metal ions [5-7]. General anesthetics have been shown to interact in the ferritin cavity within an interhelical dimerization interface[8].

Metal ion binding sites are observed at several places on ferritin: the iron entry channel, the ferroxidase center, and the nucleation site. Besides these conventional sites, an additional metal binding site was reported between the iron entry channel and the ferroxidase site of ferritin [9,10]. Others have reported that the presence of divalent cations, such as cadmium (Cd), can improve the crystallization of ferritin [11], presumably through electrostatic forces of attraction. Moreover, it has been reported that changes in the electromagnetic fields, temperature, and pH can all influence the binding of metal ions with proteins, particularly ferritins as they transport Fe ions [12-20].

## 2. Materials and Methods

### Crystallization, data collection and structure determination

Crystals of ferritin (Sigma F4503 Lot# SLBV4372) were obtained using the microbatch method under oil[21] at room temperature (293 K). The microbatch plates (Hampton Research HR3-121) wells were prepared with 2 μl CdSO_4_ in 2% agarose pellets. Equine Spleen Ferritin (Sigma F4503) without further purification was dissolved in buffer (3.5mM NH4SO4 and 100mMTris at pH7.4); 2 μl were dispensed on top of the CdSO_4_ agarose pellet and covered with 6 μl of mineral oil. Plates were submitted to electric field for 70 hours in the instrument described in Rubin and coworkers [20]. Number of crystals decreased while crystal sizes increased (**Figure 1**). After removal of the field, crystals were kept until flash-cooled directly in a nitrogen stream at 100 K.

**Figure 1.**
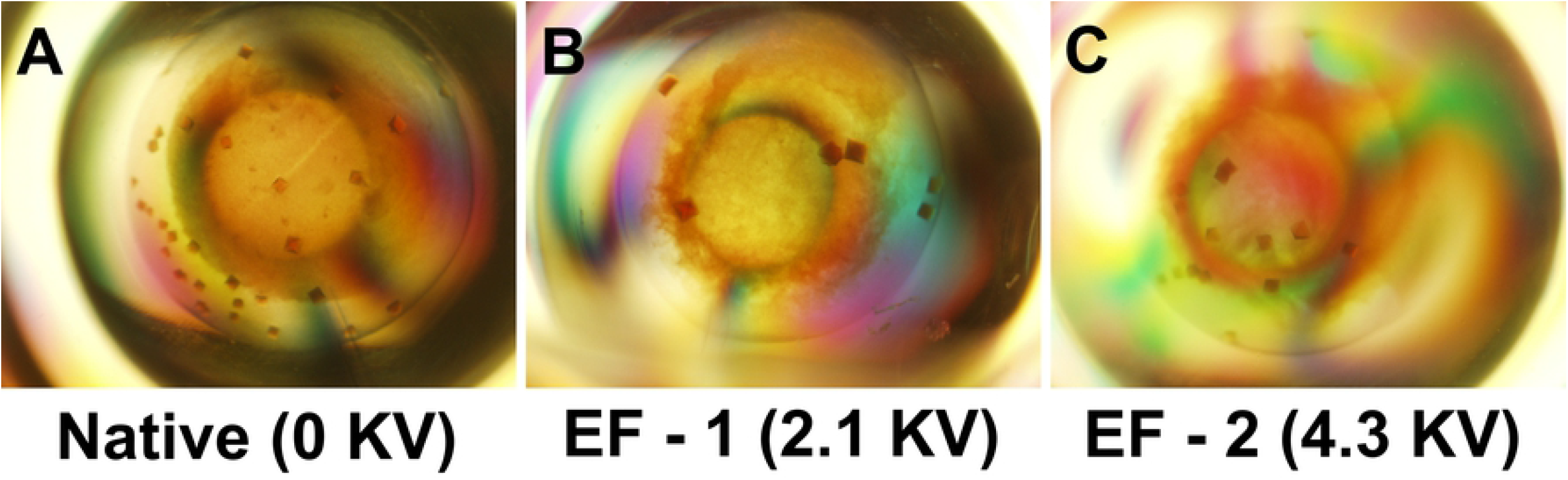
Ferritin crystals grown by the microbatch method [21] in the presence of agarose and electric field. (A) Native (B) EF-1, 2.1 KV and (C) EF-2, 4.3 KV. As the electric field intensity increases, the number of nucleation sites decreases while the size of the crystals increased.

X-ray diffraction data were collected on 19ID NYX beamline, NSLS-II. Diffraction data were processed using XDS [22]. The structures were determined by molecular replacement method with *Dimple* of ccp4i package [23] using the PDB 2W0O as search model. Structure model building and refinement were performed with *Coot* [24] and Phenix [25] software respectively. The refined model contains one monomer in the asymmetric unit with residues 2–171. All structural figures were prepared using *PyMOL* (http://www.pymol.org/2/).

### PDB deposition

Atomic coordinates and structure factors of native, EF-1 structure and EF-2 structure of ferritin have been deposited in the Protein Data Bank with code 7RYW, 7RZN and 7RZX respectively.

## 3. Results and Discussion

Ferritin crystals were grown by the microbatch under oil crystallization method [21] in the presence and absence of an external electric field using similar experimental setup described by Rubin *et al* at room temperature (293 K) [20]. In brief, the CdSO_4_ dissolved in agarose gel was added to the micro batch plate prior to the addition of the crystallization buffer and the protein solution; the crystallization drops were covered with oil to allow for controlled equilibration of the solution. The plates were exposed to electric field for 70 hours to allow for nucleation and crystallization. Notably, the number of nucleation centres were reduced with the increased electric field environment. Thus, the number of crystals that were grown decreased with the intensity of the electric field while the size of the crystals was increased (**Figure 1**).

Crystals were harvested and cryo-cooled without further cryo protectant in a nitrogen stream kept at 100 K; except for the crystal grown in the presence of 4.3 KV which was cryo-protected with 15% glycerol. One crystal for each electric field, 0 KV, 2.1 KV and 4.3 KV was exposed to X-rays, and these crystals diffracted to 2.09, 1.97 and 2.44 Å resolution, respectively. The structures were solved by molecular replacement method **(Table 1)**. Hereafter, the structures obtained from crystallization in the presence of 2.1- and 4.3-KV **<u>e</u>**lectric **<u>f</u>**ield are referred as EF-1 and EF-2 structures, respectively, with the structure obtained in the absence of an electric field referred to as the *native* structure. The structures were well defined in the electron density maps, with no residues in the disallowed regions of the Ramachandran plot. Structural analysis shows that the ferritin molecule consists of five helices and a long inter-helix loop at the N-terminus, which is located on the exterior surface of the molecule **(Figure 2)**. The fifth helix (helix α5) runs from the outside of the molecule to the inside, with the C-terminus protruding into the inner surface of the molecule. All three structures superimposed well, with no significant conformational changes observed with the changing electric field conditions (rmsd < 0.3 Å for all Cα atoms), with the exception of double conformations that were observed at the position His49 only in the native structure, in which His49 is coordinating with a Cd ion (site-5).

**Table 1.** Crystallographic data collection and refinement statistics of ferritin.

| **Table 1**<br>Data-collection and structure-refinement statistics |  |  |  |
| --- | --- | --- | --- |
| Crystals | Native | EF-1 (2.1 KV) | EF-2 (4.3 KV) |
| Data collection |  |  |  |
| Beamline | 19-ID, NSLS-2 | 19-ID, NSLS-2 | 19-ID, NSLS-2 |
| Wavelength (Å) | 0.979 | 0.979 | 0.979 |
| Space group | *F*4<sub>3</sub>2 | *F*4<sub>3</sub>2 | *F*4<sub>3</sub>2 |
| *a*, *b*, *c* (Å) | 181.73,181.73,181.73 | 182.13,182.13,182.13 | 181.07, 181.07,181.07 |
| Resolution range (Å) | 28.70–2.09 (2.15-2.09) | 28.80-1.97 (2.02-1.97) | 28.63-2.44(2.51-2.44) |
| Unique reflections | 15632(1090) | 18877(1341) | 9949(704) |
| Multiplicity | 51.7(51.3) | 23.1(24.3) | 23.4(22.2) |
| Completeness (%) | 99.8 (98.1) | 99.8(97.1) | 99.8(98.5) |
| Average *I*/σ(*I*) | 42.1 (6.60) | 31.50(4.3) | 16.0(2.5) |
| Average CC<sub>1/2</sub> | 1.0 (0.954) | 0.999(0.940) | 0.998 (0.766) |
| *R*<sub>merge</sub> | 0.112(0.815) | 0.096(0.782) | 0.244(1.323) |
| dI/sig(dI) | 0.935 | 0.904 | 0.828 |
| Structure refinement |  |  |  |
| No. of reflections | 14816 | 9426 | 17183 |
| Completeness (%) | 99.88 | 99.65 | 95.70 |
| *R*<sub>work</sub> (95% of data) | 0.16456 | 0.2128 | 0.1909 |
| *R*<sub>free</sub> (5% of data) | 0. 19267 | 2605 | 0.2529 |
| R.m.s.d., bond lengths (Å) | 0.012 | 0.001 | 0.001 |
| R.m.s.d., bond angles (°) | 1.66 | 1.307 | 1.397 |
| No. of atoms |  |  |  |
| Protein | 1370 | 1363 | 1363 |
| Metal Ions | 14 | 14 | 14 |
| Water | 100 | 205 | 64 |
| Ramachandran favored (%) | 93.0 | 93.6 | 93 |
| Ramachandran allowed (%) | 7.0 | 5.8 | 7.0 |
| Ramachandran outliers (%) | 0 | 0 | 0 |
| Clashscore | 2.17 | 2.4 | 2.6 |
| PDB Code | 7RYW | 7RZN | 7RZX |

Values in parentheses are for the outermost resolution shell.

**Figure 2.**
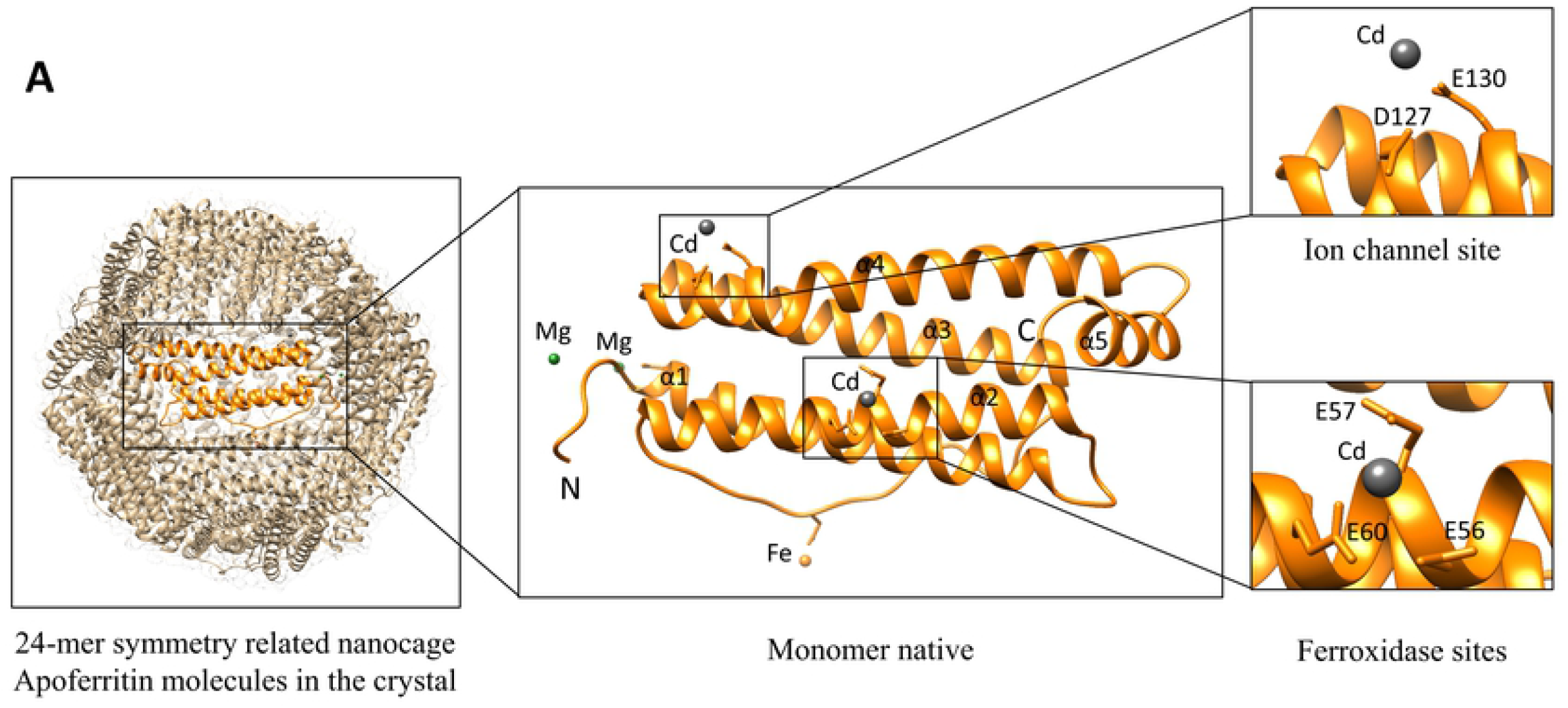
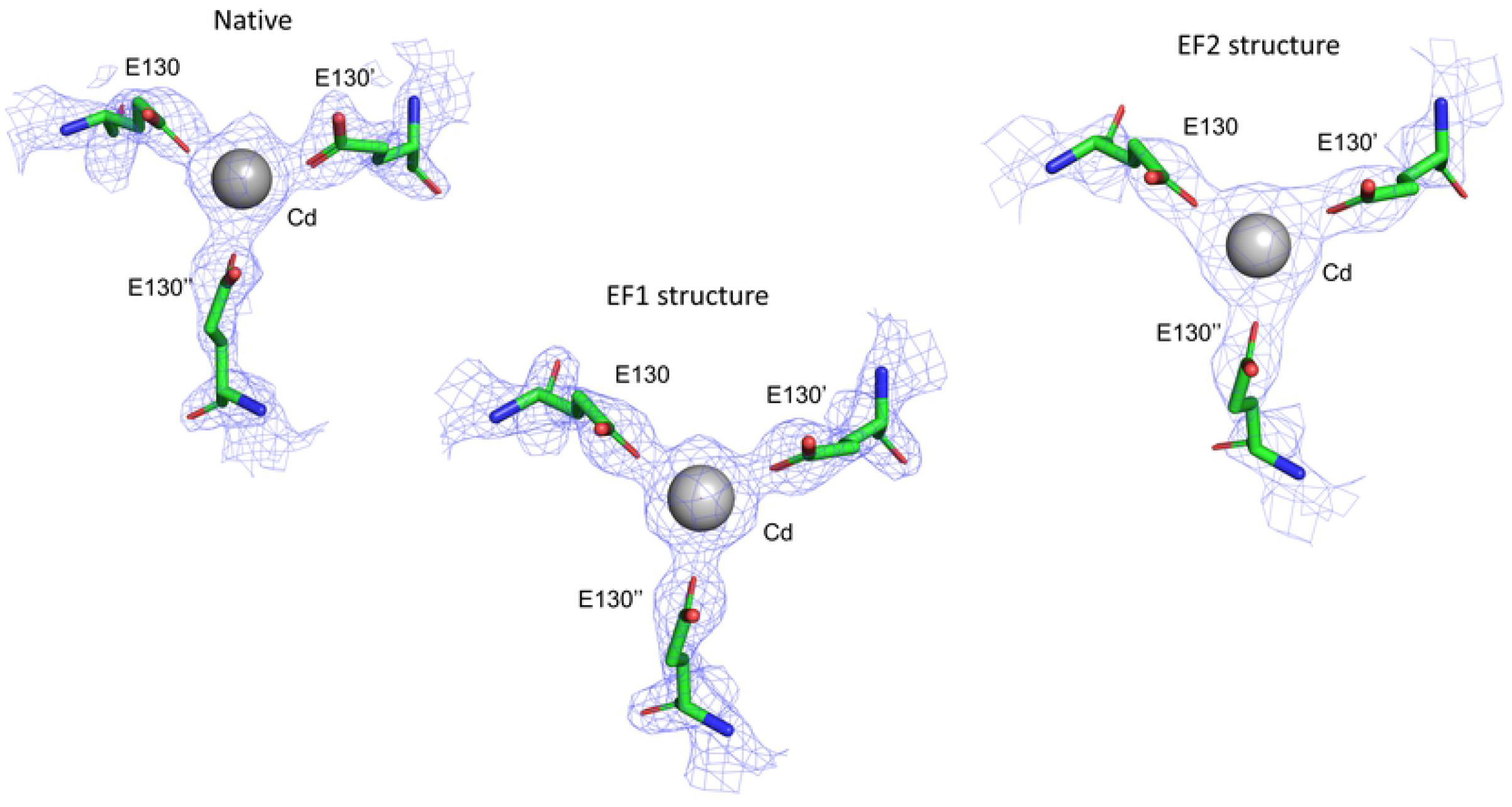
Crystal structure of ferritin. **A)** The cartoon representation of ferritin 24-mer symmetry related nanocage and the monomer showing the location of key metal ion binding sites. The helix, N and C – terminals are labelled. Close-up view of the ion channel (site 10) and ferroxidase (site 6) sites with bound metal ion are included. The atomic coordinates of the native structure were used to generate these figures. **B)** The representative *2Fo-Fc* final electron density map shows the metal ion coordination site for native (0 KV), EF-1 (2.1 KV) and EF-2 (4.3 KV) structures. The coordination of metal ion and residues sidechains, Asp127 and Glu130, are shown. The maps are contoured at 1 α level. The Cd ions are shown as grey spheres. The figures were prepared using PyMOL[26].

Notably, in these three structures, regardless of the electric field, we observed over 10 metal ions bound sites with ferritin (**Table 2**). Further these ions were mostly located within the interior of the 24-mer nanocage, which was generated by symmetry-related molecules of ferritin. In the following paragraphs, we compare the various metal binding sites in order of their relevance among several apoferritin structures.

**Table 2.**
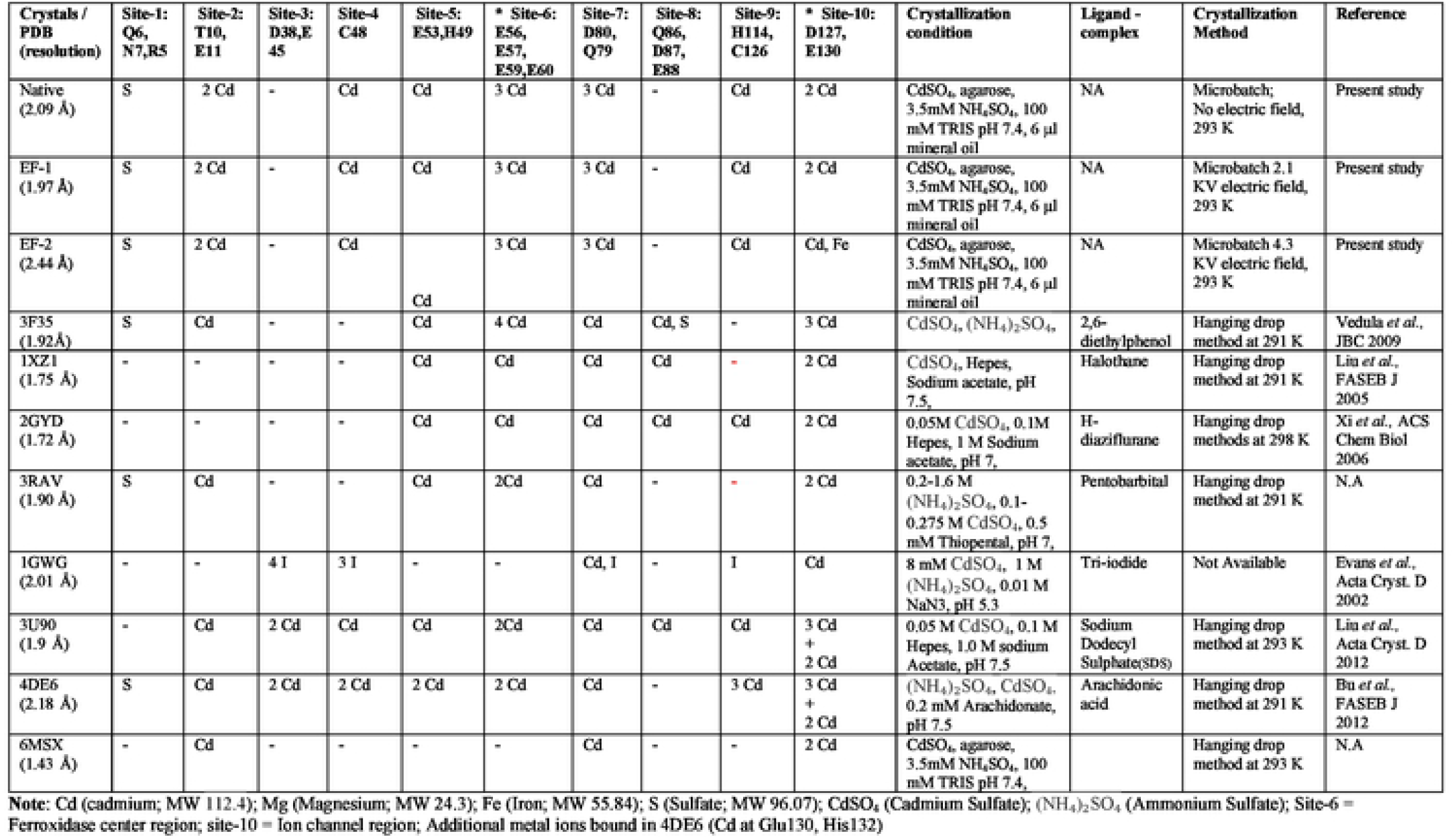
Comparison of metal ion binding sites of ferritins crystallized in different conditions.

Table 2 shows a comparison of representative ferritin structures from the PDB database reported in complex with metal ions and small molecules. These structures have the same sequence and the same basic crystallization buffers. The cation coordinating residues are mainly Glu and Asp, with some exceptions. Among the 10 binding sites compared, the ion channel region (site-10 at Asp127, Glu130) is a strong cationic binding site. In all these structures, we observed two or more Cd ions in this position (except in 1GWG with 1 Cd ion). In our three structures we consistently observed 2 Cd ions in this position as compared to 1 to 3 Cd ions reported in other structures. Similarly, we observe 1-2 Cd ions at the ferroxidase center region (site-6 at Glu56, Glu57, Glu59 and Glu60) for most of the structures except for the structure 3F35 which presents 4 Cd ions. Notably 3 Cd ions were observed in this site in our structures, which is overall a higher number than the other structures reported in the literature (**Table 2**). These observations suggest that site-6 and site-10 are strong cation interaction sites of ferritin.

Next, site-7 (at Asp80) is occupied by a Cd ion in all structures compared (**Table 2**); notably in our native, EF-1 and EF-2 structures, this position is occupied by three Cd ions, while in the other structures only 1 Cd ion is observed. A similar cation binding property is observed for site-2 (at Glu11). These might be due to the differences in the crystallization methods and conditions.

Interestingly site-1 (Arg5, Gln6 and Asn7) is differently occupied by anionic sulfate in most of these structures suggesting that this site might be an anion binding site of ferritin. Finally, site-5 (at Glu53) is occupied by a Cd ion in all the compared structures except in 4DE6 with 2 Cd ions (**Table 2**).

Taken together in sites 2, 6 and 7 we observed an increased number of Cd ions than the other ferritin structures compared (**Table 2**). Besides, sites 4 and 9 in our structures we consistently observed Cd ions, while no metal ions occupied these sites in several of the structures compared (**Table 2**). Further we observed in our structures that the highly conserved site-10 (ion channel region) is occupied by same or a smaller number of Cd ions than the other structures. Overall, our observations indicate that an increased number of ion bindings in the divalent cation binding sites in our structures. Thus, our novel microbatch agarose crystallization method retained higher number of divalent metal ions to mimic the possible *in vivo* environment.

Next, a closer inspection of these metal ion binding sites highlighted subtle differences in the orientation of these metal ion coordinating sidechains. Yet, the overall structure of the ferritin molecules was conserved, as were the metal ion coordinating residues at the ion channel region and ferroxidase center (site-10 and site-6, respectively) among the homologs.

None of the interacting metal ions (e.g., cation Cd) including at the conserved binding pockets (e.g., site-6, ferroxidase centre region; and site-10, ion channel region) of the protein were affected by the presence of the electric field. The cations Cd and Fe naturally prefer to bind to the negatively charged binding pocket (e.g., site-6 and -10) through electrostatic forces of attraction, with the interactions between the metal ions and the metal binding sites of the protein likely dominated by attractive van der Waals forces and electrostatic interactions (**Figure 2**).

Interestingly, most of structures compared in **Table 2** show Cd as the predominant cation occupying these binding sites. Indeed, CdSO_4_ is one of the main compounds in these crystallization buffers. Notably, a higher number of negatively charged residues were found at strongly conserved cationic binding sites, such as site-6 (ferroxidase center) and site-10 (ion channel region). In several of these structures, more than one cation occupied the cationic site, with there being a preference for Cd for coordination. We surmise that when a crystallization buffer without Fe is used, these positions would be occupied by an excess of available Cd ions. Overall, this comparison of metal ion binding sites and the number of metal ions bound suggests that there is no change in the metal ion binding property in response to the application of an electric field. On the other hand, our microbatch agarose crystallization method retained more number of metal ions with the protein. It is tempting to propose that this microbatch agarose crystallisation method will facilitate to retain more bound metals ions with the proteins.

In conclusion, here we report for the first time the effect of microbatch agarose crystallization of ferritin with and without an electric field. A comparison of our ferritin crystal structures with the ferritin structures reported by others using same or different crystallization conditions has revealed the presence of increased number of bound ions in our ferritin structures. There are no significant changes on the structure of ferritin irrespective of the increased metal ion bindings. We observed that the externally applied electric field during crystallization decreased the number of nucleation and thus the size of the crystals increased. Irrespective of the external electric field, this microbatch agarose crystallization method retains higher number of bound metal ions with ferritin to mimic the possible *in vivo* environment.

## Author Contributions

V.S. designed the research. A.S., P.R. and M.I. performed the research. All authors contributed to data analysis and helped writing the paper.

## Acknowledgement

This research used resources of the CBMS center supported by NIH-NIGMS #P30GM133893, and by the DOE-BER #KP1605010 and Beamline 19-ID, NYX, managed by the NYSBC, a not-for-profit consortium, at NSLS II. National Synchrotron Light Source II, a U.S. Department of Energy (DOE) Office of Science User Facility, is operated for the DOE Office of Science by Brookhaven National Laboratory under Contract No. DE-SC0012704.

